# A genome wide analysis of *Escherichia coli* responses to fosfomycin using TraDIS-Xpress reveals novel roles for phosphonate and phosphate transport systems

**DOI:** 10.1101/2019.12.23.887760

**Authors:** Muhammad Yasir, Keith Turner, Sarah Bastkowski, Ian Charles, Mark A. Webber

## Abstract

Fosfomycin is an antibiotic which has seen a revival in use due to its unique mechanism of action and resulting efficacy against isolates resistant to many other antibiotics. Mechanisms of resistance have been elucidated and loss of function mutations within the genes encoding the sugar importers, GlpT and UhpT are commonly selected for by fosfomycin exposure in *E. coli*. There has however not been a genome wide analysis of the basis for fosfomycin sensitivity reported to date. Here we used ‘TraDIS-Xpress’ a high-density transposon mutagenesis approach to assay the role of all genes in *E. coli* in fosfomycin sensitivity. The data confirmed known mechanisms of action and resistance as well as identifying a set of novel loci involved in fosfomycin sensitivity. The assay was able to identify sub domains within genes of importance and also revealed essential genes with roles in fosfomycin sensitivity based on expression changes. Novel genes identified included those involved in glucose metabolism, the phosphonate import and breakdown system, *phnC-M* and the phosphate importer, *pstSACB*. The impact of these genes in fosfomycin sensitivity was validated by measuring the susceptibility of defined inactivation mutants. This work reveals a wider set of genes contribute to fosfomycin sensitivity including core sugar metabolism genes and two transport systems previously unrecognised as having a role in fosfomycin sensitivity. The work also suggests new routes by which drugs with a phosphonate moiety may be transported across the inner membrane of Gram-negative bacteria.

**Importance:** The emergence and spread of antibiotic resistant bacteria had resulted in increased use of alternative drugs which retain efficacy against isolates resistant to other classes of drugs. One example is fosfomycin; an old drug which has found greatly increased use in recent years. We studied the mechanisms of fosfomycin resistance by applying a genome wide screen based on comparing the fitness of a massive library of transposon mutants in the presence of fosfomycin. This approach identified the previously known mechanisms of resistance but also identified a number of new pathways which contribute to fosfomycin sensitivity including two importer systems. This information advances our knowledge about an increasingly important antibiotic and identifies new potential routes to resistance.

## Introduction

The increasing prevalence of bacteria which are resistant to clinically important antibiotics has led to searches for alternative options to treat problematic infections (1). There has been limited progress in the development of new antibiotics and one strategy has been to revive older drugs which may be effective but are not common in clinical practice (2). One example is fosfomycin which has seen a sharp increase in clinical use in recent years. Fosfomycin has a unique mode of action where it targets the initial stages of peptidoglycan biosynthesis by acting as a phosphoenolypyruvate analogue and inhibiting MurA (3). This means that fosfomycin retains activity against strains producing beta-lactamases as it targets an earlier stage in peptidoglycan biosynthesis. This is attractive given the high prevalence of production of beta-lactamase enzymes of many families in important pathogens. Fosfomycin is a phosphonic-acid molecule produced in nature by *Streptomyces* species and is commonly used for treatment of complicated urinary tract infections and increasingly for more serious systemic infections. In *Enterobacteriaceae*, fosfomycin enters the cell by acting as a mimic for two nutrient importer systems; GlpT and UhpT (4).

Resistance to fosfomycin has been shown to be relatively easy to select *in vitro* and resistant mutants often show loss of function of GlpT or UhpT, or have mutations in adenylcyclase (*cyaA*), or the phosphotransferase (*ptsI*) both of which control intracellular levels of cyclic-AMP, which in turn regulates expression of *glpT* and *uhpT* (5–9). In addition, alterations within the drug target MurA (particularly those altering a Cys115 residue near the active site) can decrease susceptibility by reducing its affinity for fosfomycin (10–12). Over-expression of *murA* has also been observed in resistant isolates and is thought to confer decreased susceptibility by saturating the drug (13, 14). Finally, horizontally acquired enzymes (FosA and FosB) that inactivate fosfomycin by breaking its oxirane ring can cause resistance (15–17).

Whilst it has been easy to select for fosfomycin resistance isolates *in vitro*, there is evidence that selection of resistance carries a major fitness cost, and that fosfomycin resistant isolates may be compromised in virulence (18–20). Various studies have looked for the prevalence of fosfomycin resistance in different settings and, in general resistance rates have remained relatively low even in high use settings (21, 22).

Given the recent increase in the use of fosfomycin (23) and the lack of clarity around aspects of its mode of action and resistance we used a genome wide transposon mutagenesis approach in *E. coli* to identify loci involved in fosfomycin susceptibility. We identified new loci as being involved in fosfomycin sensitivity including the phosphonate uptake and utilisation system, phosphate import system as well as identifying sub domains within some genes involved in fosfomycin sensitivity.

## Results

### Susceptibility of BW25113 to fosfomycin

Baseline susceptibility to BW25113 was measured and the MIC determined to be 4µg/ml, the transposon mutant library was then inoculated into fresh media (~10^7^ mutants were added to each reaction) containing multiples of the MIC and allowed to grow overnight. Replicates were completed with and without the presence of IPTG as an inducer (at two different concentrations) of the outward facing promoter present in the transposon to give a total of 30 independent experiments. After incubation, DNA was extracted from all reactions and transposon insert sites identified by sequencing. Supplementary Figure 1 shows the concordance between independent repeats and the impacts of increasing drug concentrations on numbers of reads mapping across the genome.

### Genes involved in fosfomycin susceptibility

Whilst different drug concentrations did identify some specific loci as important in survival there was a clear core set of genes identified as being important in all conditions (Supplementary Figure 2). A total of 31 separate loci were identified as being involved in determining sensitivity to fosfomycin at all exposure concentrations (Table 1). These included the majority of known, chromosomal mechanisms of resistance with strong signals identified for *murA*; the target for fosfomycin, *glpT* and *cyaA*. This validates the specificity of TraDIS-Xpress in identifying genes involved in fosfomycin susceptibility. The TraDIS-Xpress method also proved able to assay essential genes. For example, *murA* is an essential gene but mutants with inserts which are positioned upstream and in the same orientation as *murA* were highly enriched in the presence of IPTG (Figure 1). These mutants will over-express *murA* which will help the target saturate the activity of fosfomycin. The high density of the library also allows very high resolution, for example *cyaA* was identified as a significant target but enrichment of inserts was restricted to this gene’s regulatory domain (Figure 1). Mutations within this domain have previously been reported as being important to determine fosfomycin sensitivity (but not the rest of the protein) and TraDIS-Xpress was able to clearly identify the sub-domain within the gene responsible for fosfomycin. Interestingly, no signal was identified at the *uhpT* locus which has also previously been implicated in fosfomycin import although is not expressed in the test conditions so would not be under selective pressure.

**Figure 1.**
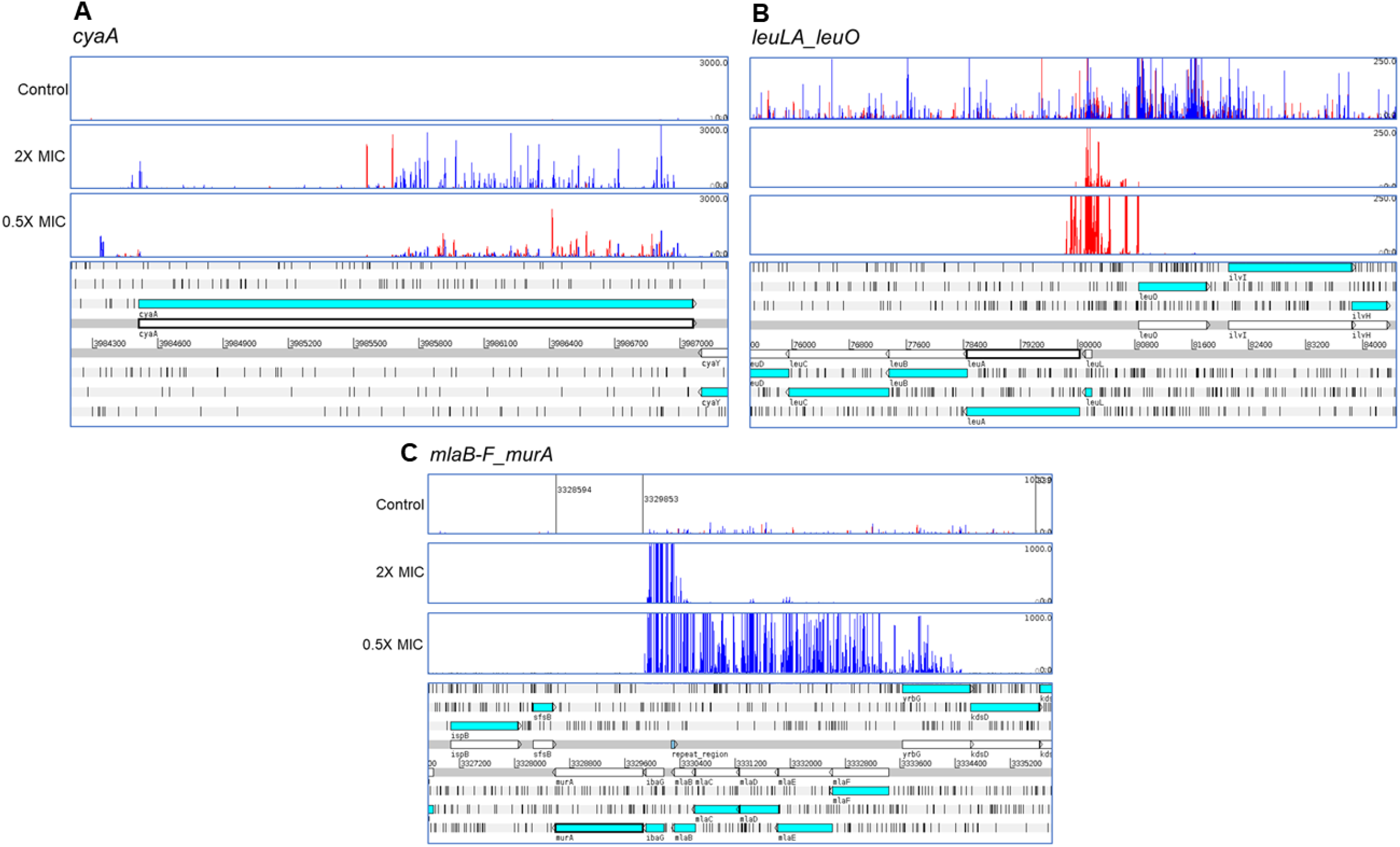
Differential selection of transposon mutants at *cyaA* (**A**), *leuO* (**B**) and *murA* (**C**). The bottom of each panel illustrates the genomic context and the panels above illustrate the mapped reads, red bars indicate reads orientated left-to-right and blue bars the opposite. The height of each bar reflects abundance of each insert.

**Table 1.** Loci significantly altered in all exposure conditions.

| **Locus** | **Predicted impacts of insertion** | **Function and interpretation** |
| --- | --- | --- |
| *ahr* | Over-expression | Aldehyde reductase |
| *alsR* | Inactivation and up-regulation of *alsBACEK* | Allose binding transporter |
| *cmk* | Inactivated | Cytidylate kinase; nucleotide binding |
| *cra* | Inactivated | Catabolite repressor/activator. Global regulator of carbon metabolism |
| *crp* | Antisense knockdown | Global regulator and carbon catabolite repression |
| *cyaA* | Inactivation of regulatory domain | Adenylate cyclase; regulates cyclase activity by carbon source |
| *dacB* | Intragenic insert at one site | Peptidoglycan biosynthesis |
| *galU* | Inactivated | Synthesis of UDP-D-glucose |
| *glpE* | Inactivation and overexpression of *glpD* | Glycerol metabolism |
| *glpG* | Inactivated | Membrane protein protease |
| *glpK* | Inactivated | Glycerol kinase; key player in glucose control of glycerol metabolism |
| *glpT* | Inactivated | Glycerol uptake; known fosfomycin importer |
| *glpX* | Protected | Fructose bisphosphatase |
| *guaA* | Inactivation | GMP synthesis |
| *ibaG* | Inactivated, inserts upstream of *murA* | Increase of *murA* expression |
| *leuA* | Protected and over-expression of *leuO* | Regulation of stringent response, pleiotropic effects from *leuO* |
| *leuL* | Inactivated and over-expression of *leuO* | Pleiotropic effects from *leuO* |
| *mlaB-F* | Inactivated – inserts upstream of *murA* | Peptidoglycan recycling; fosfomycin target expression increased |
| *murA* | Upstream inserts | Peptidoglycan biosynthesis; fosfomycin target; expression increased |
| *mutL* | Protected | DNA repair |
| *nagB* | Inactivated with up-regulation of *nagE* | Over-expression of sugar importer (*nagE*) |
| *phnC-M* | Inactivated (all 14 members of operon in sense) | Phase variable phosphonate transporter and degradation complex |
| *phoU* | Inactivation (downstream of *pstB*) | Repressor of *pstABCS* phosphate uptake system |
| *pstABCS* | Inactivated | Phosphate uptake system |
| *ptsH* | Inactivated | Decoration of imported sugars with phosphates |
| *purA* | Inactivated | Purine biosynthesis |
| *rfaH* | Inactivated (inserts antisense to *tatD*) | Transcription anti-terminator |
| *tatD* | Protected | DNA repair |
| *treC* | Inactivated | Hydrolyses trehalose to glucose |
| *waaFGP* | Inactivated | LPS branching |
| *yhfA* | Protected and up-regulated | Conserved hypothetical product, known to be regulated by CRP |

### Identification of new mechanisms of fosfomycin resistance

As well as loci known to be involved in determining fosfomycin sensitivity several new loci were identified by Tradis-Xpress. These included several genes involved in sugar metabolism beyond those already noted as contributing to fosfomycin susceptibility (table 1). Significant patterns of inserts were seen at *cra, crp, cyaA, galU, glpK, glpX* and *treC*. Together these genes are all involved in control of available glucose within the cell which will in turn influence expression of the glucose importers that are known routes of entry for fosfomycin. Mutants that over-expressed *leuO*, a pleiotropic regulator with known roles in control of stress responses were strongly selected by fosfomycin (figure 1). As were mutants with inserts within the phosphonate and phosphate uptake systems (figure 2). The *phnC-*P phosponate uptake and degradation system is a large operon composed of an ABC importer (*phnCDE*), regulator (*phnF*), a multisubunit phosphonate degradation complex (*phnG-L*) and some ancillary components (*phnM-P*). The system has been reported to be phase variable due to the presence of either 2 or 3 copies of a repeat element within *phnE* and the downstream genes (PhnF-L) are thought to be cryptic in K-12. Despite this, there was a massive enrichment of inserts throughout this system upon exposure to fosfomycin (figure 2), sequencing of the parent strain of the library, and mapping of from reads obtained after fosfomycin exposure confirmed our strain carried 2/3 inserts and was, therefore supposedly cryptic. However, the TraDIS-Xpress data suggested a large fitness advantage from inactivation of most of this system. Additionally, another ABC importer (PstSACB) was also implicated in fosfomycin sensitivity (figure 2) with inactivation of this system favouring growth in the presence of fosfomycin. Both *mutL* and *tatD*, involved in DNA repair processes were protected under all fosfomycin exposure conditions.

**Figure 2.**
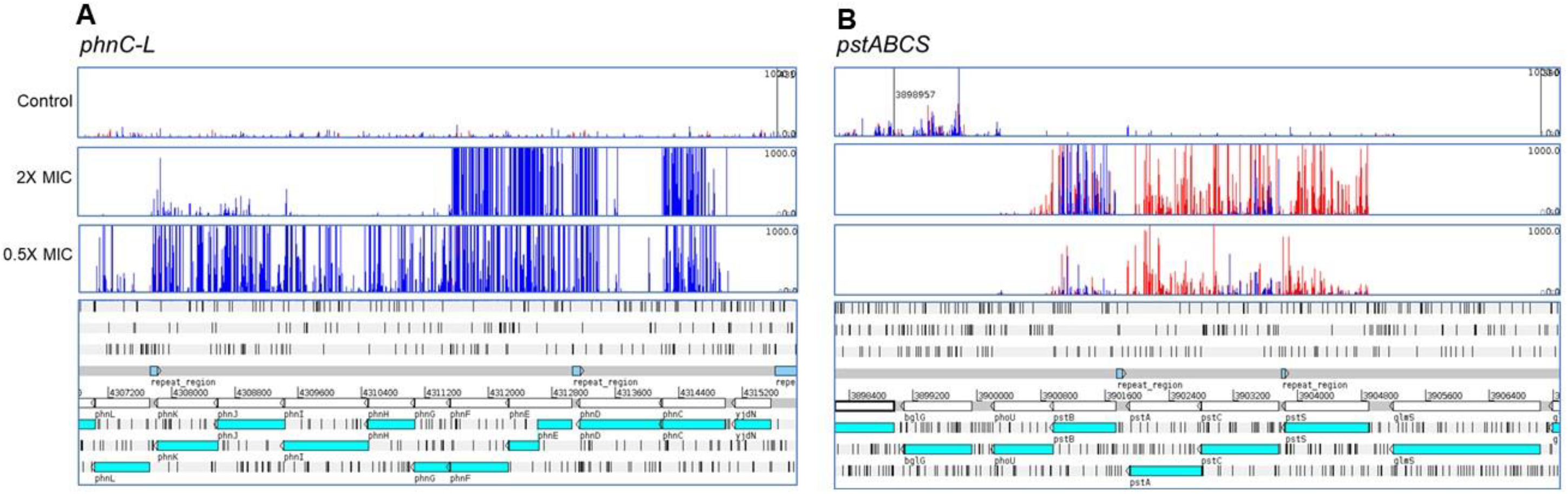
Inactivation of the *phn* (**A**) and *pst* (**B**) operons is selected by fosfomycin. The bottom of each panel illustrates the genomic context and the panels above illustrate the mapped reads, red bars indicate reads orientated left-to-right and blue bars the opposite. The height of each bar reflects abundance of each insert.

None of the genes important in all conditions have been associated with resistance to other drugs, i.e. there was no signal observed for porins or multidrug efflux pumps or their regulators suggesting selection of cross-resistance to other agents by fosfomycin may be limited.

### Validation of targets

To test the predictions made by TraDIS-Xpress we selected a set of nine genes and took both the corresponding mutants from the KEIO collection for each and tested their sensitivity to fosfomycin by growth on agar containing different concentrations of the drug (Figure 3). BW25113 was inhibited by the MIC of the drug as expected, as were both *leuO* mutants tested; this was as expected as the data predicted *leuO* over-expression was important for survival. For all the other mutants one or both mutants tested showed significantly improved growth compared to the parent strain. This was particularly evident for mutants in three separate parts of the *phn* operon or the two *pts* mutants tested (Figure 3). Given the indicated role for the phosphonate uptake system we also tested whether addition of an exogenous phosphonate would impact fosfomycin sensitivity. We used etidronate, a small bis-phosphonate and found in checkerboard assays with etidronate and fosfomycin a consistent rescue of growth to 1-2 dilutions above the MIC of fosfomycin by addition of 400 mg/L of etidronate.

**Figure 3.**
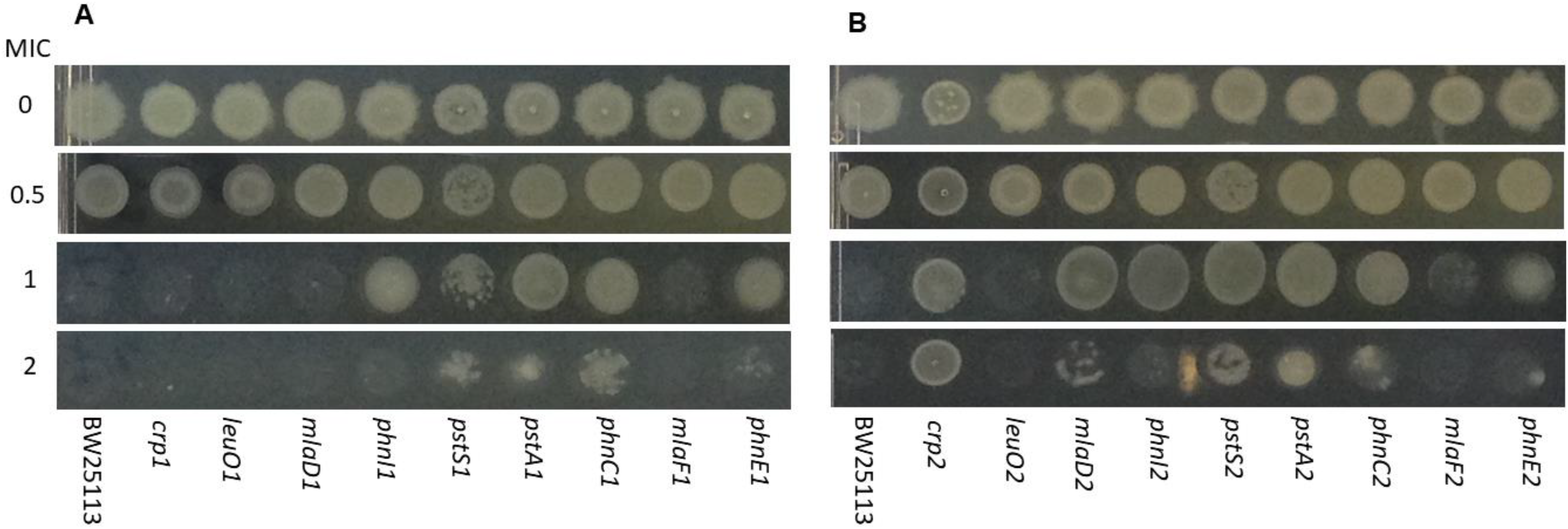
Validation of specific mutants by inoculation onto agar containing different fosfomycin concentrations. Panels **A** and **B** represent analysis of both independent mutants for each gene present in the KEIO collection. For each strain, 5µl spots representing ~10^4^ cfu were inoculated and incubated overnight at 37 °C.

## Discussion

The mechanisms of fosfomycin action and resistance known to date have largely been elucidated by the study of individual mutants and isolates that demonstrate fosfomycin resistance (18). Here, we used TraDIS-Xpress to assay the role of the whole genome of *E. coli* on fosfomycin sensitivity on one experiment. The data showed the sensitivity of this approach with all the known, chromosomal mechanisms of resistance being identified as important contributors to fosfomycin susceptibility (apart from *uhpT* which is not expressed in the assay conditions used). Additionally, TraDIS-Xpress was also able to identify domains within genes which are important, for example the regulatory domain of CyaA is known to be involved in fosfomycin susceptibility by controlling cAMP levels and therefore expression of *glpT* and *uhpT* (19). Inactivation of the domain of *cyaA* responsible for this results in lower expression of both importers and decreased susceptibility to fosfomycin. TraDIS-Xpress clearly indicated the role for this regulatory domain but not the rest of *cyaA* (figure 1).

Traditional TraDIS experiments have not been able to assay essential genes which are often the targets for antibiotics, TraDIS-Xpress was able to clearly show a key role for MurA in fosfomycin sensitivity with very strong enrichment of mutants that were positioned upstream of this gene and in the same orientation – these will over-express *murA* helping to saturate fosfomycin and allow escape of its inhibition of peptidoglycan biosynthesis.

In addition to genes known to be involved in fosfomycin resistance a wider set of new loci were identified as contribution to sensitivity to the drug (table 1). This included a set of genes involved in glucose metabolism, again likely to be mediating expression of the glucose importers which facilitate transport of fosfomycin across the inner membrane. This extends our current understanding of how central metabolism can impact sensitivity to fosfomycin and shows how specific growth conditions (and associated expression of genes) are likely to dictate sensitivity to this drug by altering gene regulation. Genes involved in DNA repair (*mutL* and *tatD*) were protected in the presence of fosfomycin, this may represent damage to DNA resulting from altered metabolism following inhibition of the primary target and consequent bactericidal effect. This has been proposed as a common impact of many bactericidal drugs (24–26).

Whilst UhpT and GlpT are known import systems for fosfomycin, mutation of either the PhnC-M or PstBCSA systems was strongly selected for by fosfomycin exposure (figure 2). The phosphonate uptake and metabolism system is made up of a 14 gene operon containing an ABC transporter, phosphonate lyase complex (able to break the P-C phosphonate bond) and regulatory genes. Fosfomycin contains a phosphonate moiety and so is a plausible substrate for this system. The TraDIS-Xpress data showed a very strong signal for enrichment of mutants inactivating components of this system with selection for inactivation of all genes at sub-MIC exposures and the ABC transporter and regulator above the MIC (figure 2). This suggests inactivation of the ABC importer provides a strong fitness advantage in the presence of fosfomycin suggesting this is another, unrecognised route of import for the drug. This system has been suggested to be cryptic in K-12 with expression being recovered after a phase variation mediated alteration of the copy number of a repeat element within *phnE*. This is however downstream of the start of the operon with the first components of the ABC transporter upstream of this site so their expression would not be influenced by this site in any case. The TraDIS-Xpress data does strongly suggest a biological benefit from inactivation of most genes in the operon at some fosfomycin conditions, this challenges the notion that this system is cryptic in these conditions in K-12. Mutants in three parts of the operon all demonstrated a growth benefit in the presence of fosfomycin, supporting a role for inactivation of the system. Addition of the bis-phosphonate, etidronate did result in limited (1-2 fold MIC increase) rescue of growth in the presence of fosfomycin – this data is supportive of the phosphonate importer being an additional route of entry into the cell for fosfomycin.

The potential for phosphonate moieties to mediate import of molecules into the cell is intriguing and may have potential utility, a major challenge in therapy is getting drugs into cells and new routes to modify molecules and promote their uptake are likely to enhance efficacy of drugs. It has become clear in recent years that most drugs cross the inner membrane by active import rather than passive diffusion (27) and identifying side groups which may improve uptake by changing importer specificity is important.

Similarly, the phosphate uptake system, PstBCAS was also identified as being involved in fosfomycin sensitivity with inactivation of the system proving beneficial in the presence of the drug (figure 2). These data suggest that multiple importers, including novel systems are involved in fosfomycin sensitivity, testing of defined mutants in both the Phn and Pst systems confirmed a phenotype with the mutants able to grow above the MIC of the drug (figure 3).

One major worry with some drugs is selection of cross-resistance to other agents, this is often mediated by generic mechanisms of resistance including multidrug efflux and or porins where expression changes can influence accumulation of many drugs (28). There was not a strong signal for these pathways after fosfomycin exposure which suggest the major mechanisms of fosfomycin resistance (figure4) are likely to be relatively specific and not influence other classes of agent. This is potentially important as, whilst fosfomycin resistance is not hard to select a key feature of this drug is its activity against strains resistant to other drugs, in particular various beta-lactams.

**Figure 4.**
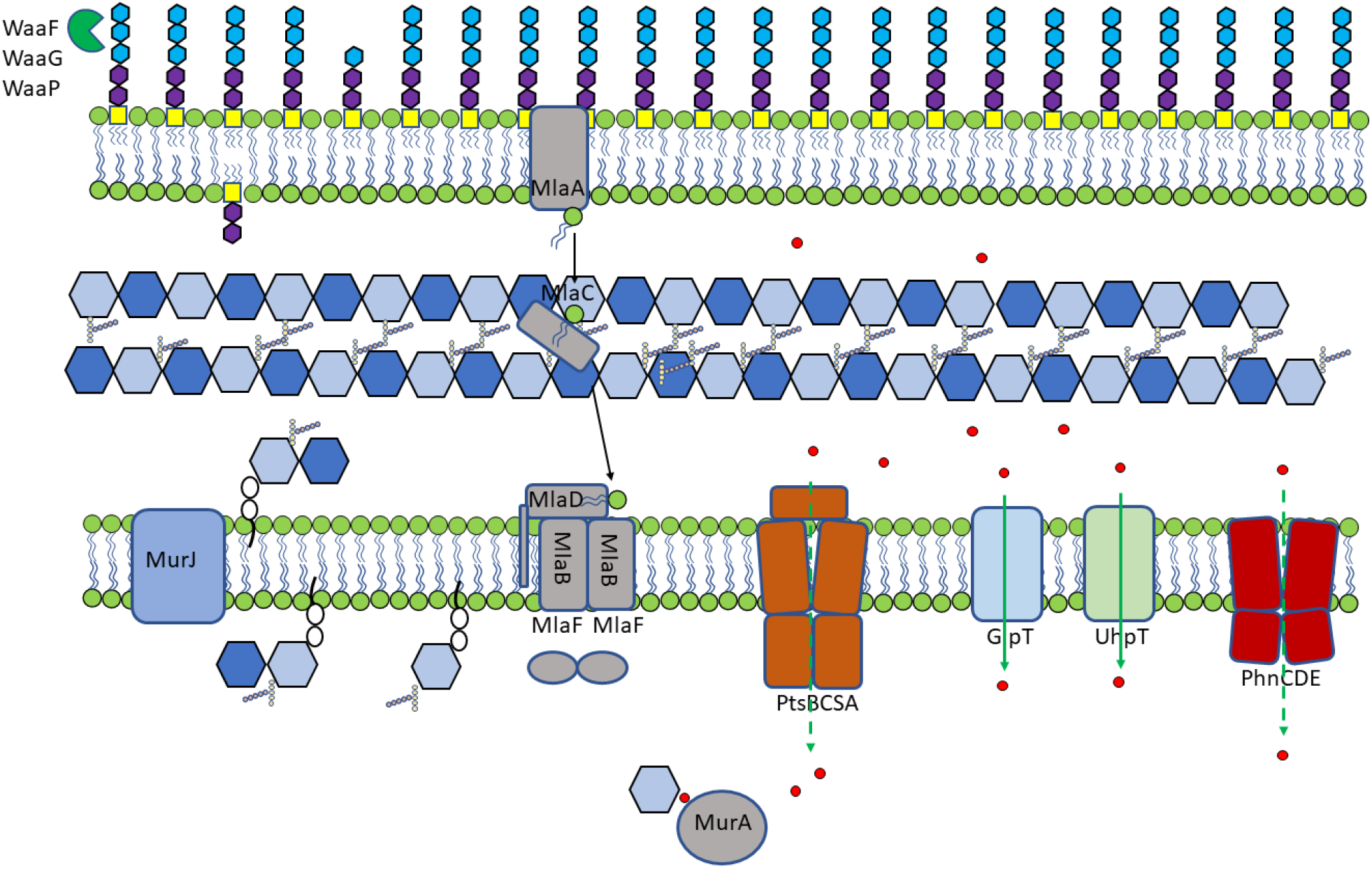
Genes identified as being involved in fosfomycin sensitivity. Fosfomycin is indicated by red circles and known and potentially novel mechanisms of uptake are indicated by filled and dashed green arrows, respectively.

Taken together, these data show a genome-wide analysis of genes involved in sensitivity to fosfomycin identifies new loci which are important for determining sensitivity to the drug, suggest important roles for two transport systems not previously implicated in fosfomycin resistance as well as providing new information about potential routes for drug entry into cells.

## Materials and Methods

### TraDIS-Xpress library

We recently described the construction of a high density TraDIS-Xpress library in *E. coli* BW25113 which was used in this work (29). The transposons (a mini-Tn5 transposon coding for kanamycin resistance (*aph(3’)-Ia*)) used contain an outward facing *tac* promoter 3’ to the kanamycin cassette which is inducible by IPTG allowing over-expression or repression of genes (depending on insert orientation) as well as traditional inactivation. This allows the roles of essential genes in a stress to be analysed based on expression changes; traditionally these loci have been cryptic in TraDIS experiments as insertions within them are lethal.

### Fosfomycin exposure conditions and TraDIS-Xpress sequencing

The minimum inhibitory concentration (MIC) of fosfomycin against BW25113 was determined using microbroth dilution in LB broth, the same growth medium that was used for TraDIS-Xpress experiments. For TraDIS-Xpress experiments, approximately 10^7^ mutants were inoculated into LB broth containing doubling concentrations of fosfomycin ranging from 0.25X MIC to 2X MIC (and a drug free control). Replicate experiments were completed with no induction, or with the addition of 0.2 or 1 mM IPTG to induce transcription from outward oriented promoter. Mutants were allowed to grow for 24 hours at 37°C. All experiments were performed in duplicate to give a total of 30 independent TraDIS-Xpress experiments.

After growth in experimental conditions, DNA was extracted from pools of mutants using a ‘Quick-DNA™’ Fungal/Bacterial 96 kit extraction kit (Zymo Research). DNA was then fragmented using a Nextera DNA library preparation kit (Illumina) with the modification of Tnp-i5 oligonucleotides being used instead of i5 index primers, and 28 PCR cycles. The resulting DNA was size selected to purify fragments between 300bp-500bp and sequenced on a NEXTSeq 500 sequencing machine using a NextSeq 500/550 High Output v2 kit (75 cycles).

### Bioinformatics

Results were analysed using the AlbaTraDIS tool (version 0.0.5) which we developed for TraDIS-Xpress analysis and recently described (29). Briefly, sequence reads were mapped against the BW25113 reference genome (CP009273), aligned and insertion plots created.

The patterns of inserts were compared between fosfomycin exposed and control conditions, AlbaTraDIS calculated the number of inserts within each gene as well as assessing the number of ‘forward’ and ‘reverse’ insertions per gene and, within a window of 198 bp upstream and downstream of each gene. The number of sequence reads are modelled on a per-gene basis using a negative binomial distribution and an adapted exact test as implemented in edgeR (30) followed by multiple testing correction (31) to identify significant differences between conditions. A set of default cut-offs for significance and number of reads were applied (q-value <= 0.05, logFC >= 1, logCPM > 8). The resulting gene list provided a prediction of genes involved in fosfomycin survival as well as a prediction whether a change in expression of a gene (either down- or up-regulated) influences survival. The insertion patterns at candidate loci were visually inspected using ‘Artemis’ which was also used to capture images for figures. (32)

### Validation experiments

A total of 18 mutants were selected from the KEIO library to validate predictions about sensitivity to fosfomycin made from by TraDIS-Xpress (33). These included genes in the phosphonate uptake and metabolism and phosphorous import systems identified as major contributors to fosfomycin sensitivity as well as a set of randomly selected control genes not expected to have any impact on fosfomycin sensitivity. Both replicate mutants present in the KEIO collection for each gene were independently analysed. Mutants were tested for their sensitivity to fosfomycin by MIC determination. All experiments were duplicated (giving at least four datasets for each gene; two repeats from each of the two mutant alleles of each gene). BW25113 was included in all experiments as a control.

## Data access

All sequence data have been deposited with EBI under project accession number PRJEB29311.

